# Phenotypic link between protein efficiency and pig welfare suggests no apparent trade-offs for mitigating nitrogen pollution

**DOI:** 10.1101/2023.03.03.530955

**Authors:** Lea Roch, Esther Oluwada Ewaoluwagbemiga, Claudia Kasper

## Abstract

Pig manure contributes significantly to environmental pollution through nitrogen compounds. Reducing protein in feed can help, but it may lead to damaging behaviors if pigs’ nutritional needs are not met. Breeding pigs for higher protein efficiency (PE) is a long-term solution to reduce nitrogen pollution, but concerns about pig welfare remain. We studied 95 pigs involved in a project on the genetic basis of PE on a 20% protein restricted diet to investigate the phenotypic connection between PE and welfare. These pigs represented natural PE variations in the population. At around 100 days, before their PE was known, we observed their behaviors. Only three pigs engaged in tail biting and manipulation of vulnerable regions, but this was not associated with PE. There was no clear link between PE and manipulating pen mates’ less vulnerable regions. Such behaviors are normal but can cause stress and injury if carried out excessively due to boredom or stress. Overall, pigs with higher PE showed no major behavioral abnormalities in this study. Considering the lack of genetic knowledge, the risk of increased harmful behaviors when selecting for higher PE appears low when inferred from this purely phenotypic association.

## Introduction

Pig production contributes to environmental pollution: applying pig manure to fields releases nitrogen compounds, leading to eutrophication of soils and water bodies^1, 2^. Especially in areas with high livestock density and low availability of arable land, this problem has become particularly severe. Furthermore, the low self-sufficiency of protein sources for pig feed of the EU results in the need to import soybean meal for animal feed, mainly from South America, USA and China. This exposes the European pig industry to an increased risk of sustainability problems due to differing production standards, long-distance transport and land-use changes in regions where soy is grown on deforested land^3^, as well as shortages and price volatility of soybeans on the world market^3, 4^. Increased efforts are being made across Europe to limit pigs’ protein intake to mitigate the environmental damage from pig production. Pomar and Remus^5^ showed that each percentage reduction in dietary nitrogen leads to a 1.5% decrease in nitrogen excretion. However, van der Meer *et al*.^6^ reported an increase in damaging behaviors, including tail and ear biting, during a 20% reduction in pigs’ dietary protein, especially when combined with poor sanitary conditions. In a trial where a 20% reduction in protein was combined with restricted feeding and an unbalanced supply of essential amino acids (EAA), especially methionine, an outbreak of tail biting occurred after two months, which subsided when access to feed was increased^7^. Conceivably, deficiencies in certain essential AAs, such as methionine, threonine, and tryptophan, result in neurotransmitter system dysfunction, as specific AAs serve as neurotransmitters or are required for neurotransmitter synthesis^6^. Moreover, foraging behavior could increase due to the relative scarcity of AAs, potentially leading to obsessive manipulation and biting behaviors, especially when straw is unavailable as enrichment^8^. Further, AA deficiency is thought to make blood taste more attractive^9^. The trade-off between ensuring animal welfare and rendering pig production more environmentally friendly is apparent: while restricting the protein in feed can potentially reduce nitrogen emissions, it is important to examine whether this measure will affect pigs’ wellbeing.

Damaging behaviors, including tail biting, are common in pig production, resulting in compromised victim welfare and economic losses. Tail biting and excessive manipulation also indicate that a pig’s behavioral needs are unmet, suggesting poor wellbeing of the biters^8^. Outbreaks of tail biting are often difficult to identify, have a multifactorial origin and occur when stressors accumulate, including a lack of suitable occupational materials, poor climate conditions in the barn, inadequate cleanliness, unbalanced diet and poor health. Behavioral changes at the pen level, such as in feeding patterns, can be detected up to one month before a tail biting outbreak occurs^10^. At an individual level, tail posture has also been shown to be an early pen-level indicator of an impending outbreak of tail biting^11^ and behavioral problems caused by various stressors begin before escalating into harmful behavior leading to serious injury. For instance, a pig could proceed from gently manipulating a conspecific’s tail or ear, with no noticeable reaction from the target, to biting and inducing a bloody wound^6, 12, 13^. Ursinus *et al*.^14^ found a positive correlation between the number of penmate manipulations, a generally natural and harmless behavior, and a high rate of tail biting. There is a delicate boundary between natural pig behaviors, including exploring and manipulating conspecifics, and damaging behaviors, such as biting. For instance, engaging in positive social behaviors has a positive impact on growth, presumably by reducing stress through oxytocin release^13^. However, redirected “abnormal” behaviors, such as excessive belly nosing, ear biting, and “tail-in-mouth”^15^, can severely disturb the receiver^16^, cause permanent stress and even impact growth^13^.

Individual differences in protein efficiency (PE), i.e., the ability to utilize dietary proteins^17^, are heritable^18, 19^. Therefore, it is to be expected that individual pigs will differ in how their behavior is affected by a protein reduction. Harnessing these heritable individual differences in breeding would therefore effectively reduce the long-term nitrogen pollution from pig production. The molecular basis of PE is not yet well established, and there are legitimate concerns that breeding for increased PE could induce behavioral problems and reduced welfare. For instance, Breuer *et al*.^20^ reported a weak but significant genetic correlation between the lean tissue growth rate and tail biting behavior in Landrace pigs, but not in Large White pigs, the breed used in the present study. Whether PE or related traits have any connection with the likelihood of becoming a victim of damaging or problematic behaviors is not yet known. Therefore, there seems to be a certain risk of inadvertently co-selecting pigs with an increased predisposition to behavioral problems when breeding for a higher protein accretion rate. However, the aim of sustainable pig production should not be to increase the rate of protein accretion, but rather to increase PE, as this takes into account not only the output in terms of muscle mass, but also the input in terms of the amount of protein consumed. PE is a trait arising from a combination of several processes, including gastrointestinal tract absorption and protein turnover, which occur in different tissues and organs. Genetic selection for higher PE could improve these processes or could alter the allocation of proteins toward lean tissue growth and away from other processes, including immune or endocrine system functioning, potentially compromising homeostasis and thus health and reproduction. The genetic architecture of PE or nitrogen excretion is likely complex, as multiple regions on different chromosomes were found associated with nitrogen excretion traits^21^. Some of these quantitative trait loci (QTLs) overlap with various production traits, while others are unique to nitrogen excretion^21^, though the functions of genes within these QTLs are still unclear.

In this exploratory study, we aimed to investigate whether PE is associated with indicators of impaired welfare on a phenotypic level in pigs subjected to dietary protein restriction. Particularly, whether the level of PE is associated with performing or receiving tail biting and other damaging behaviors was examined, as were the performed and received manipulation behaviors of pen mates. Furthermore, we explored the relationship between PE and straw rooting, which is considered an indicator for positive welfare, allowing animals to explore the environment, thereby reducing boredom and manipulation of pen mates^16^. Here, we also investigated the outcome of social encounters and confrontations related to PE as an indicator of whether aggressiveness or dominance is associated with efficiency. Finally, we explored the relationship of the frequency of lesions, tail position, cleanliness, and activity with PE.

## Material and Methods

### Animals and diets

The pigs were part of a larger experiment comprising 681 pigs in 14 farrowing series (batches) of the Swiss Large White dam line herd at the experimental farm Agroscope Posieux, with the goal of estimating genetic parameters of PE^19^. The pigs were not selected for increased PE. In this study, 95 non-tail-docked pigs (53 females and 42 castrated males) born in two farrowing series were observed between August and December 2020. One male pig died before slaughter, so PE could not be assessed. About seven days after birth, male piglets were surgically castrated. The piglet was removed from the box and an anti-inflammatory drug (ketoprofen) was injected. After 15 min, the piglet was anaesthetized with isoflurane inhalation for 90 s and the testes were removed. The piglet was placed in a warm cradle until it woke up (3-4 min after the end of the isoflurane inhalation) and, when fully awake, was returned to the box with the sow and littermates. The pigs were raised in 36.78 m^2^ pens with 23 or 24 pigs in each, i.e., two pens per series. The floor space was above legal requirements, with only partial slatted flooring (as required by law in Switzerland). Pens were cleaned daily, and the pig had ad libitum access to drinking water and a low-crude-protein diet, which was distributed by single-spaced automatic feeding stations (Schauer Maschinenfabrik GmbH & Co. KG, Prambachkirchen, Austria), which and were accessible to the pigs via individual identification by an RFID chip in the ear from 07:40 to 23:30, allowing individual feed intake monitoring. Pigs were fed a standard starter feed after weaning until reaching an average body weight of 22.4 (± 1.6) kg. They were then mixed in groups of 24 pigs, in which they stayed until slaughter, and received a grower diet that was 20% protein-reduced compared to the recommended diet in Switzerland^22^. From an average body weight of 63.5 (± 2.4) kg, the pigs were fed a 20% protein reduced finisher diet. All EAA were reduced to the same level to avoid deficiencies and thus a nutritional imbalance. The chemical composition of the grower and finisher diet is shown in Table 1. Each pen was covered with a thin layer of straw on the concrete floor, contained two mobile and two fixed straw baskets, and held four suspended metal chains as occupational materials. Following the Swiss Animal Welfare Act, it was ensured that occupation material, in our case unchopped straw, was always available. The two groups had auditory and olfactory contact. Pigs had no outdoor access. Natural light was provided through windows in the barn. The health status of the pigs on each observation day was considered, and each injured animal was treated on the same day.

**Table 1:**
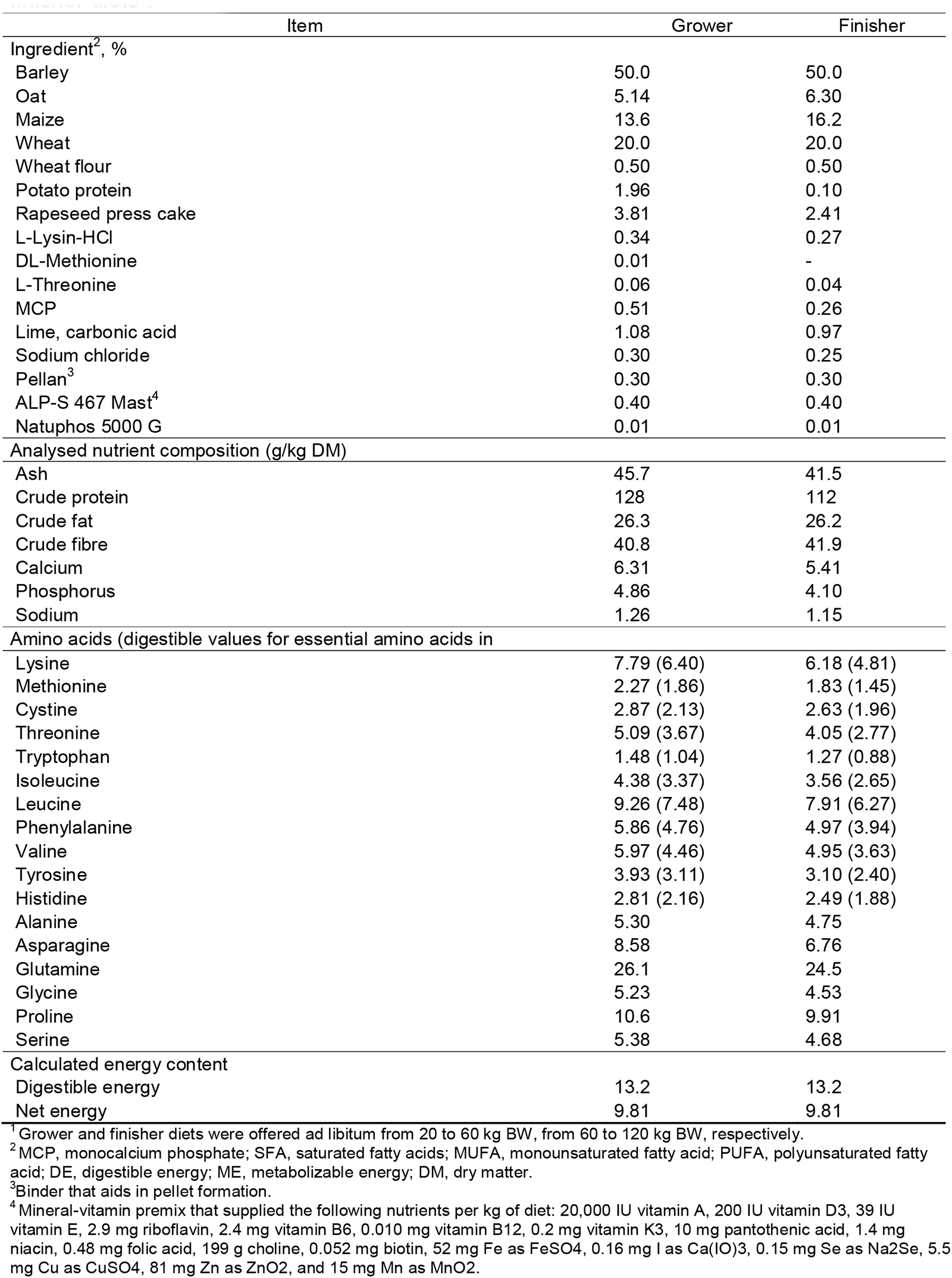
Dietary ingredients and analysed composition (g or MJ/kg as-fed) of the grower and finisher diets^1^.

### Behavioral observations

Between days 98 and 115 after birth, each pig’s behavior was recorded following an ethogram (Table 2), which had been created based on literature research and two days of observations, and refined during a preliminary study prior to this work. The first farrowing series was observed in August 2020 and the second in December 2020. Direct behavioral observations were conducted in daylight between 12:45 and 16:00. Prunier *et al*.^23^ suggested that pig activity peaks in the early morning and afternoon. As the pens were cleaned in the morning, the observations were made in the afternoon, as this was the time of least external disturbance. However, the pigs might have rested more during hot periods in the first farrowing series. The observer (LR) stood outside the pen to limit any influence on the animals. Observations began after a 15-min habituation period. All pigs in one pen were individually marked with colored spray and directly observed for 5 min using focal sampling^24^ in a random sequence on one afternoon^25^. Each pen was observed over four different days; thus, each pig was observed for 20 min total, resulting in approximately 65 h of observation, including scanning for lesions, tail position, and cleanliness, as described in Table 3. During observation, the observer was blind to the PE of the individual pigs, as this was only recorded several weeks later, at the time of slaughter. The total number of separate instances of nasal and oral manipulation of objects (metal chains, pen barriers, and drinkers), as well as of pen mates during the 5 min of individual observation, was recorded. These behaviors comprised biting, seizing (with the mouth, but not the teeth), and manipulation (with a closed mouth or the snout). Behaviors directed toward the head, body, ears, tail, and vulva or perineal area of a conspecific were recorded separately, and we recorded whether the focal animal performed (active) or received the behavior (passive). Due to the scarcity of manipulation behaviors directed at pen mates, we later combined the behaviors: all bites directed (received) toward the ears, tails, and vulva or perineal (ETV) region were summed into the variable “damaging behaviors”. All other behaviors, including biting, seizing, and manipulation with the snout, directed (received) at the ETV region, were aggregated into one variable, termed “problematic behaviors,” and biting, seizing, and manipulation with the snout directed toward the rest of the body (excluding the ETV region) into another variable (“potentially problematic behaviors”) by totaling the instances of each behavior. The same was done for behaviors received.

**Table 2:** Ethogram of observed pig behaviours (modified from Roch *et al.* ^24^)

| Category of behaviour | Description of behaviour | Target of behaviour |
| --- | --- | --- |
| Oral and nasal actions |  |  |
| Biting | Opens and closes jaw with force at least once, and uses teeth | Objects, pen mates |
| Seizing | Opens and closes the jaw without force at least once, not using teeth | Objects, pen mates |
| Manipulation | Keeps the jaw closed, manipulates with the snout, the snout is mobile. The jaw may be relaxed and mobile but the individual does not grasp with the mouth | Objects, pen mates |
| Rooting | Manipulates the straw on the ground or in a basket with the snout, often bites into the straw or eats the straw | Straw on the ground or in a basket |
| Confrontations between pen mates | Pen mates physically oppose each other with face-to-face or head-to-body contact, with a push that can be gentle to strong | Pen mates |

The start and the outcomes of confrontations, i.e., agonistic situations arising when two pigs meet and both want an object or place simultaneously, were recorded, and the following were possible: win (domination), lose (submission), or a tie, when the outcome was unclear. The number of instances of straw rooting was determined as the sum of the number of times the focal pig was engaging with the baskets and whether it performed straw rooting behaviour on the floor (never in 5 min = 0, ≥ 1 s in 5 min = 1). In addition, all wounds and scratches (classified as mild scratches or severe wounds), the tail posture (raised curled, hanging straight, or tucked between legs), and the cleanliness of each pig were recorded after each observation, using the protocol in Table 3.

**Table 3:** Observations of lesions, tail posture and cleanliness of the pigs (from Roch *et al*.^24^)

| Observation | Categories |
| --- | --- |
| Lesions<sup>1</sup> | 0: no lesion<br>1: superficial lesions (scratches)<br>2: wounds (deep lesions, clearly visible fresh or dried blood)<br>3: part of tail or ear missing |
| Quantity of lesions | few: no lesions or fewer than five<br>many: five or more lesions |
| Tail posture<sup>2</sup> | 1: curled<br>2: tail straight or hanging<br>3: tail tucked between legs |
| Cleanliness<sup>3</sup> | 1: clean (less than 10% of the body surface covered in excrements)<br>2: slightly soiled (10 to 30% of the body surface covered in excrements)<br>3: heavily soiled (more than 30% of the body surface covered in excrements) |

<sup>1</sup>Categories inspired by Smulders *et al.*<sup>26</sup>; Ursinus *et al.*<sup>14</sup>; Valros *et al.*<sup>27</sup>; Zonderland *et al.*<sup>28</sup>)
<sup>2</sup>Categories inspired by (Ursinus *et al.*<sup>14</sup>; Zonderland *et al.*<sup>29</sup>)
<sup>3</sup>Categories from KTBL 2016 (Kuratorium für Technik und Bauwesen in der Landwirtschaft, Tierschutzindikatoren: Leitfaden für die Praxis - Schwein. Darmstadt. https://www.ktbl.de/shop/produktkatalog/12631/ (retrieved on Feb 18, 2023).

### Protein efficiency and performance traits

The PE of the 94 pigs surviving until slaughter was calculated as the amount of protein in the carcass after slaughter divided by the amount of protein ingested. In this study, we worked with the naturally occurring variation in PE in pigs not selected for this trait. Pigs were slaughtered at an average live body weight of 105.2 kg (± 6. 5) at the Agroscope experimental slaughterhouse in Posieux. After 16 h of feed deprivation, the pigs were individually transported in a trolley to the research slaughterhouse (located 100 m from the barn). They were stunned with a CO_2_ stunner (87% CO_2_; Samson C1 L 803; MPS Group, Holbaek, Denmark) for 180 s and immediately exsanguinated. The intestines and viscera, as well as the hair, hooves and blood, were removed, and the carcass was cut in half. Half-carcasses were scanned on a dual energy X-ray absorptiometry (DXA) device (GE Lunar i-DXA, GE Medical Systems, Glattbrugg, Switzerland) to determine the lean meat content. The following regression equation (1), which was developed in a previous study^30^, was used to estimate the protein content from the lean meat content, determined by DXA:

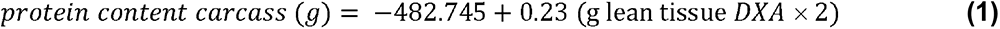

The amount of protein ingested from the start of the switch to the protein-reduced grower feed was recorded by the automated feeders. The estimated protein content of the carcass at the time of the feed change was subtracted from the total weight of protein in the carcass at slaughter (for more details see^18^). The lean and fat mass in the carcass were obtained from DXA. The information needed to calculate average daily feed intake (ADFI), average daily gain (ADG) (2) and feed conversion ratio (FCR) (3) were automatically recorded by the feeder stations and during the weekly individual weighing of the pigs.

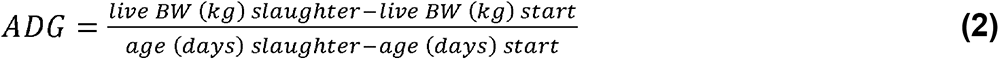

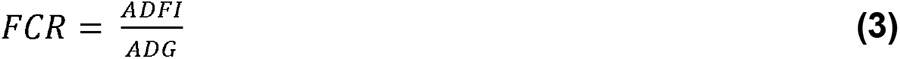

*Live BW (kg) slaughter* and *age (days) slaughter* are the live body weight in kg shortly before slaughter and the age in days at slaughter, respectively, and *live BW (kg) start* and *age (days) start* are the exact body weight in kg and the age in days at the start of the grower phase, respectively.

### Statistical analysis

We grouped the single behaviors directed toward conspecifics into the following categories with decreasing intensity and damaging capacity by calculating the total number of occurrences: (i) “damaging behaviors” – biting directed toward the ETV regions of conspecifics, which were too rare for statistical analysis; (ii) “problematic behaviors” – seizing and manipulation of conspecifics’ ETV regions; and (iii) “potentially problematic behaviors” – biting, seizing, and manipulation of all other body (non-ETV) regions of conspecifics, including the head. All different behaviors directed at objects, as well as rooting in the floor straw and in the basket, were combined into a single variable (termed “object manipulation” and “straw rooting”, respectively), and all analyses were conducted with the R software version 4.2.2^31^. We identified PE outliers using the Grubbs test in the *outliers* package version 0.15^32^. This resulted in the removal of two individuals, with a PE of 0.574 and 0.469, from further statistical analysis, leaving a data set of 92 pigs. In consideration of these data, which were count variables with likely more zeros than expected due to an important proportion of pigs not performing the behavior in question during the observation window (i.e., zero-inflated) or data that might not satisfy the strict mean–variance relationship of a Poisson distribution (i.e., over-dispersion), we used the R package *glmmTMB* version 1.1.4^33^. The effect of PE on problematic behaviors, confrontation outcomes, and straw rooting in terms of behavior counts was analyzed with generalized mixed-effects models with a Poisson, quasi-Poisson, and negative binomial family, and the latter two were chosen to account for different mean–variance relationships, i.e., over-dispersion. We included sex as a covariate in all models, and body weight at the time of observation in the models on confrontations.

Since the error distribution that would best fit each behavioral count variable was unknown at the start of the analyses, we ran several models for all variables of interest to determine the best fitting model in a model selection procedure: First, a full model with PE and sex as fixed effects was performed for each family (Poisson, quasi-Poisson and negative binomial), and the intercept was modelled as zero-inflated with a logit link. If the zero-inflation term was not significant, we reran the model without zero-inflation. If it was significant, zero-inflation was maintained for the respective error distribution models. The two nested models included one with PE only, and one with only the intercept for each model family. Thus, in total, 9 models were run. For initiating a confrontation and its outcomes, we also included body weight at the time of the observation, because it might influence the ability to perform in confrontations. Thus, we ran 12 models for confrontations initiated, won and lost: the full model including PE, sex and weight, a model with PE only, one with body weight only, and one with only the intercept. The individual ID and the pen ID nested in the farrowing group were added as random effects to correct for multiple observations and the effects of the social group on behavior, and to account for the fact that observations were carried out for two farrowing series where the temperature was rather different. Using the AICctab command from the *bbmle* package version 1.0.25^34^, the best model was chosen based on AICc (modified AIC for small sample sizes). Following Burnham and Anderson^35^, we considered models with a difference in AICc (ΔAICc) of less than two and having the same evidence, and we presented them as confidence sets. A note of caution: with this approach, a potential relationship between PE and the behaviors studied cannot be ruled out on the basis of non-significant p-values alone, i.e., p > 0.05, as it is always the case with providing conclusive evidence for the absence of an effect in the framework of null-hypothesis testing. We therefore also interpret the inclusion or absence of PE (or any other variable) in the models within the confidence sets as additional evidence. For the models in the confidence set, we used the R package *DHARMa*^36^ to diagnose any violations of distribution assumptions and model misspecifications. Plots were created using the *ggplot2* package version 3.3.6^37^ with the *ggstatsplot* extension^38^ to compute group and sex differences.

### Ethical approval

The experimental procedure was approved by the Office for Food Safety and Veterinary Affairs of the Canton of Fribourg (animal experimentation license 2018_30_FR), and all procedures were conducted in accordance with the Swiss Ordinance on Animal Protection and the Ordinance on Animal Experimentation and the ARRIVE guidelines.

## Results

### Animal performance

Some performance traits differed across farrowing series and sexes. Castrated males consumed significantly more feed and had a higher ADG than females in the second series (Table 4). Fat mass in the carcass differed between the farrowing series, with the second series having significantly lower fat mass than the first one. Within the second, but not within the first series, females had significantly lower fat mass than males (Table 4, Fig. 1). Neither the series nor the sexes differed in FCR and lean mass of the carcass (Table 4).

**Table 4:** Means and standard deviations of performance and carcass traits compared across sex and farrowing series. Significant effects (p < 0.05) are highlighted in bold.

| | group 1 | mean ± SD | group 2 | mean ± SD | t-value<sup>1</sup> | p-value<sup>2</sup> |
| --- | --- | --- | --- | --- | --- | --- |
| ADFI (kg/day) | **series 1 F** | **2.23 ± 0.23** | **series 1 M** | **2.47 ± 0.24** | **4.89** | **0.023** |
| | series 1 F | 2.23 ± 0.23 | series 2 F | 2.07 ± 0.18 | -3.98 | 0.069 |
|  | **series 1 F** | **2.23 ± 0.23** | **series 2 M** | **2.43 ± 0.19** | **4.94** | **0.022** |
|  | **series 1 M** | **2.47 ± 0.24** | **series 2 F** | **2.07 ± 0.18** | **-8.30** | **5.1$\times 10^{-5}$** |
| | series 1 M | 2.47 ± 0.24 | series 2 M | 2.43 ± 0.19 | -0.79 | 0.943 |
|  | **series 2 F** | **2.07 ± 0.18** | **series 2 M** | **2.43 ± 0.19** | **9.13** | **3.1$\times 10^{-6}$** |
| ADG (kg/day) | series 1 F | 0.83 ± 0.1 | series 1 M | 0.9 ± 0.08 | 4.01 | 0.104 |
|  | **series 1 F** | **0.83 ± 0.1** | **series 2 F** | **0.75 ± 0.08** | **-4.61** | **0.043** |
| | series 1 F | 0.83 ± 0.1 | series 2 M | 0.9 ± 0.08 | 3.96 | 0.104 |
|  | **series 1 M** | **0.9 ± 0.08** | **series 2 F** | **0.75 ± 0.08** | **-8.34** | **2.7$\times 10^{-5}$** |
| | series 1 M | 0.9 ± 0.08 | series 2 M | 0.9 ± 0.08 | -0.31 | 0.996 |
|  | **series 2 F** | **0.75 ± 0.08** | **series 2 M** | **0.9 ± 0.08** | **8.58** | **1.2$\times 10^{-5}$** |
| FCR | series 1 F | 2.7 ± 0.08 | series 1 M | 2.73 ± 0.08 | 2.10 | 1.000 |
| | series 1 F | 2.7 ± 0.08 | series 2 F | 2.78 ± 0.12 | 4.26 | 0.142 |
| | series 1 F | 2.7 ± 0.08 | series 2 M | 2.71 ± 0.08 | 0.92 | 1.000 |
| | series 1 M | 2.73 ± 0.08 | series 2 F | 2.78 ± 0.12 | 2.37 | 1.000 |
| | series 1 M | 2.73 ± 0.08 | series 2 M | 2.71 ± 0.08 | -1.19 | 1.000 |
| | series 2 F | 2.78 ± 0.12 | series 2 M | 2.71 ± 0.08 | -3.44 | 0.442 |
| lean mass (kg in | series 1 F | 62.24 ± 2.96 | series 1 M | 62.17 ± 1.9 | -0.14 | 1.000 |
| | series 1 F | 62.24 ± 2.96 | series 2 F | 60.4 ± 3.49 | -2.78 | 1.000 |
| | series 1 F | 62.24 ± 2.96 | series 2 M | 61.73 ± 1.87 | -1.08 | 1.000 |
| | series 1 M | 62.17 ± 1.9 | series 2 F | 60.4 ± 3.49 | -2.83 | 1.000 |
| | series 1 M | 62.17 ± 1.9 | series 2 M | 61.73 ± 1.87 | -1.05 | 1.000 |
| | series 2 F | 60.4 ± 3.49 | series 2 M | 61.73 ± 1.87 | 2.20 | 1.000 |
| fat mass (kg in | series 1 F | 20.72 ± 2.33 | series 1 M | 22.9 ± 3.01 | 3.74 | 0.119 |
|  | **series 1 F** | **20.72 ± 2.33** | **series 2 F** | **14.98 ± 2.39** | **-12.08** | **3.3$\times 10^{-9}$** |
| | series 1 F | 20.72 ± 2.33 | series 2 M | 19.2 ± 2.28 | -3.37 | 0.119 |
|  | **series 1 M** | **22.9 ± 3.01** | **series 2 F** | **14.98 ± 2.39** | **-12.74** | **7.4$\times 10^{-9}$** |
|  | **series 1 M** | **22.9 ± 3.01** | **series 2 M** | **19.2 ± 2.28** | **-6.14** | **0.002** |
|  | **series 2 F** | **14.98 ± 2.39** | **series 2 M** | **19.2 ± 2.28** | **8.46** | **1.1$\times 10^{-5}$** |
| PE | series 1 F | 0.38 ± 0.01 | series 1 M | 0.38 ± 0.02 | -1.26 | 1.000 |
| | series 1 F | 0.38 ± 0.01 | series 2 F | 0.4 ± 0.02 | 3.76 | 0.208 |
| | series 1 F | 0.38 ± 0.01 | series 2 M | 0.4 ± 0.02 | 4.25 | 0.129 |
| | series 1 M | 0.38 ± 0.02 | series 2 F | 0.4 ± 0.02 | 3.81 | 0.208 |
| | series 1 M | 0.38 ± 0.02 | series 2 M | 0.4 ± 0.02 | 3.98 | 0.208 |
| | series 2 F | 0.4 ± 0.02 | series 2 M | 0.4 ± 0.02 | -0.20 | 1.000 |

<sup>1</sup>Games-Howell pairwise post-hoc tests for unequal variances
<sup>2</sup>Holm-adjusted p-values
ADFI average daily feed intake, ADG average daily gain, FCR feed conversion ratio, PE protein efficiency

**Fig. 1:**
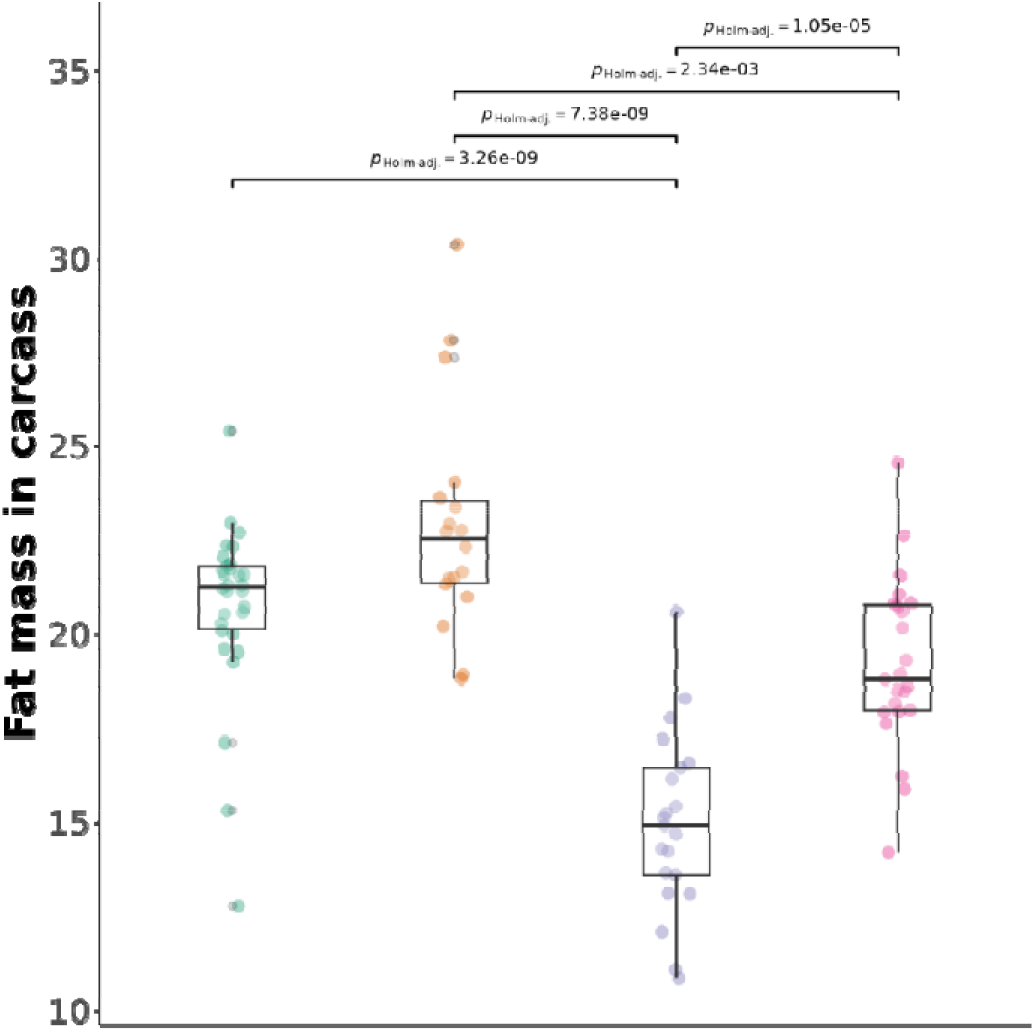
Comparison of fat mass between farrowing series and sexes using 2-sided Games-Howell pairwise tests. P-values are Holm-adjusted and bars are shown only for significant comparisons. From left to right: green: females in first farrowing series, yellow: males in first farrowing series, blue: females in second farrowing series, pink: males in second farrowing series.

### Protein efficiency

The average PE was 0.39 ± 0.02, but the farrowing series differed significantly in their PE (Fig. 2, left; Welch two-sample t-test, t = -4.03, df = 89.87, p < 0.001). The first farrowing series had a mean PE of 0.38 ± 0.02 and the second a mean PE of 0.40 ± 0.02. To account for this difference, we included the pen ID nested in farrowing series (in addition to the individual ID) in the following models. The sexes did not differ significantly in terms of PE (Fig. 2, right; Welch two-sample t-test, t = 0.20, df = 78.28, p = 0.84).

**Fig. 2:**
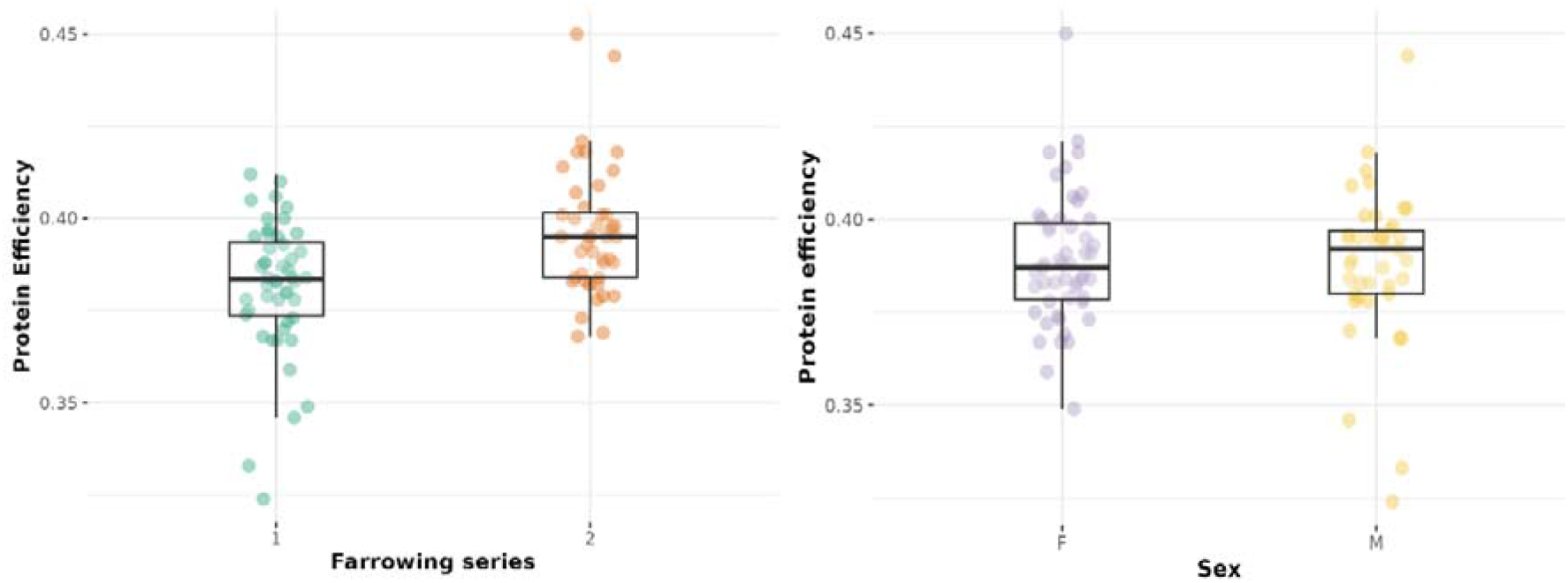
Protein efficiency (proportion of protein ingested that was retained in the carcass) in the two farrowing series (left) and sexes (right). Abbreviations: F females, M castrated males.

### Damaging and potentially problematic behaviors

Of the total 5,479 actions recorded over the four observations, 11% were directed at chains, barriers or drinkers, 27% at straw, and 61% at conspecifics. Most pig-directed behaviors targeted the body and head, i.e., less vulnerable regions, and fewer actions were directed at vulnerable regions (ETV regions; Fig. 3). Damaging behaviors, i.e., biting directed at a pen mate’s EVT regions, was observed on only nine occasions. The eight pigs performing these damaging behaviors all had an average or below-average PE. Of all 92 pigs, only three were involved in tail biting (3.2%): two bit pen mates and one was both a biter and a victim. Most pigs performed potentially problematic behaviors, but 16 pigs (17%) did not exhibit these behaviors in any of the observations. Only four pigs (4%) were not subjected to these behaviors, and only one pig was not involved at all (neither as performer nor receiver).

**Fig. 3:**
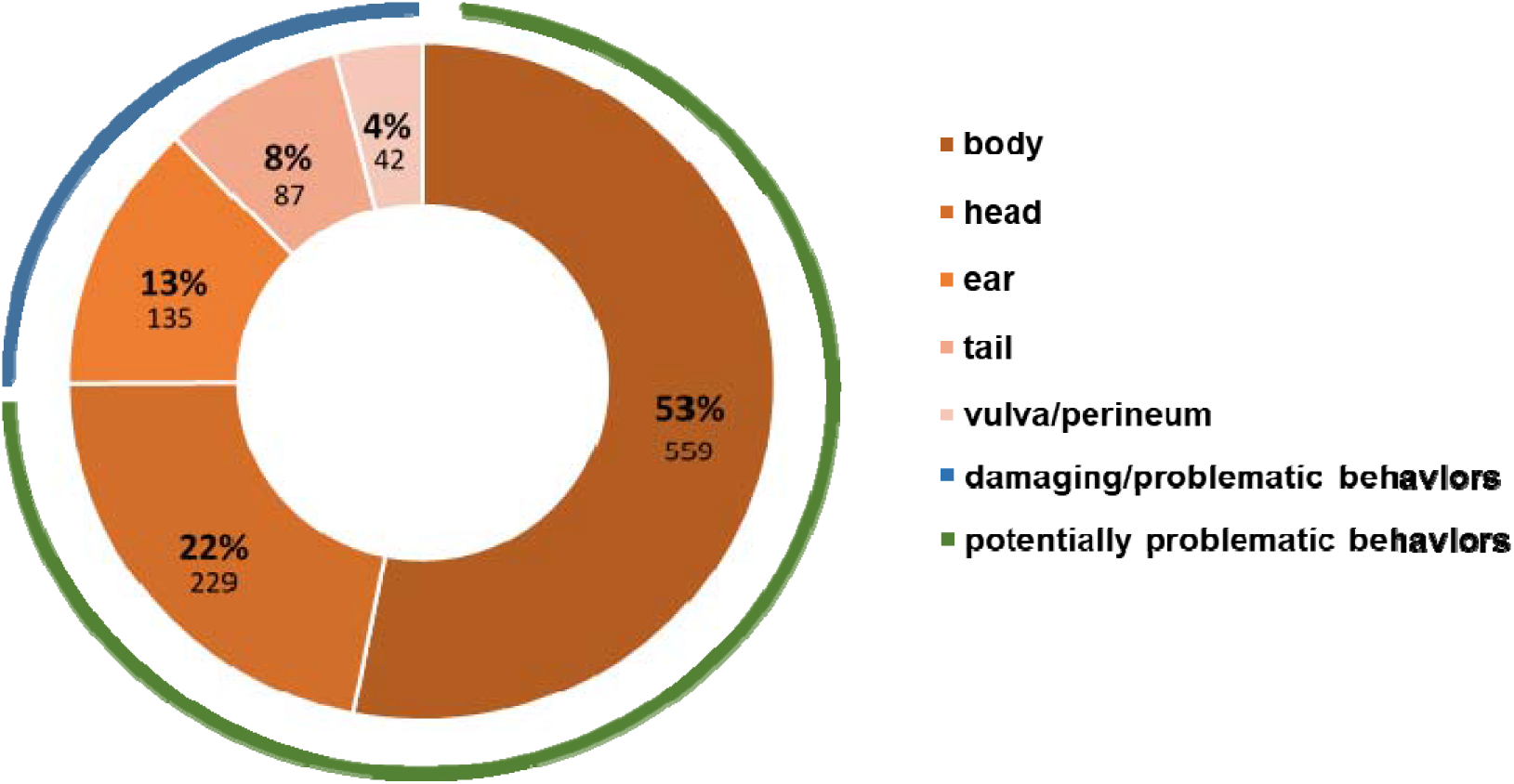
The percentages of behaviors directed at the body, head, ears, tails, vulva or perineum of all active pig-directed behaviors recorded during all observations. Absolute values are shown below in smaller print. The grouping of the behaviors into damaging (biting directed toward ear, tail, vulva or perineum) or problematic behaviors (manipulation with mouth or nose except for biting towards these vulnerable regions) is illustrated with a blue line, and potentially problematic behaviors (biting and manipulation with mouth or nose to body and head) is illustrated with a green line.

We did not find evidence that the number of problematic behaviors (seizing and manipulation directed toward a conspecific’s ETV regions) was associated with PE. In best model, PE was present but not significantly associated with the behavior (Table 5; Fig. 4). Sex was not included in the best model. There were four other models with a similar fit in terms of AICc. PE was included in two of them, but it was never significantly associated with problematic behaviors (Table S1). Neither PE nor sex was included in the best model of the number of potentially problematic behaviors (biting, seizing, and manipulation with the snout) directed at a conspecific’s less vulnerable body areas (non-ETV regions) (Table 5; Fig. 4). Another model had a similar fit in terms of AICc, and it included PE, but it was not significant (Table S1). The best model of problematic behaviors received toward ETV regions only included the intercept; thus, there was no evidence that PE or sex was associated with these behaviors (Table 5; Fig. 4). There was no other model in the confidence set (Table S2). Concerning manipulations of the whole body received, PE was included in the best model, but it was not significantly associated with the behaviors (Table 5; Fig. 4). Also in the other two equally fitting models, PE was either not included or not significant (Table S2).

**Table 5:** Best models from the model selection for problematic (ETV) and potentially problematic (head and body) behaviours performed and received as a function of protein efficiency and sex. Only the best fitting models in terms of AICc are shown.

| behavior | model | variable | estimate | SE | z-value | p-value | df | set |
| --- | --- | --- | --- | --- | --- | --- | --- | --- |
| performed ETV | QP | Intercept | -4.61 | 2.33 | -1.98 | 0.048 | 6 | 5 |
|  |  | PE | 9.31 | 5.95 | 1.57 | 0.117 |  |  |
| performed head and body | ZI-NB | Intercept | 0.89 | 0.11 | 8.15 | <0.001 | 6 | 2 |
| received ETV | ZI-PO | Intercept | -0.16 | 0.27 | -0.60 | 0.546 | 5 | 1 |
| received head and body | NB | Intercept | 3.01 | 1.89 | 1.60 | 0.110 | 6 | 3 |
|  |  | PE | -7.90 | 4.85 | -1.63 | 0.103 |  |  |

Abbreviations: ETV behaviours directed at ears, tails or perineal region of conspecifics, ZI zero-inflated, NB negative binomial, QP quasipoisson, PO poisson, PE protein efficiency, df degrees of freedom, set size of confidence set, i.e., number of models with equal fit ( $\Delta AICc \leq 2$ to the best model)

**Fig. 4:**
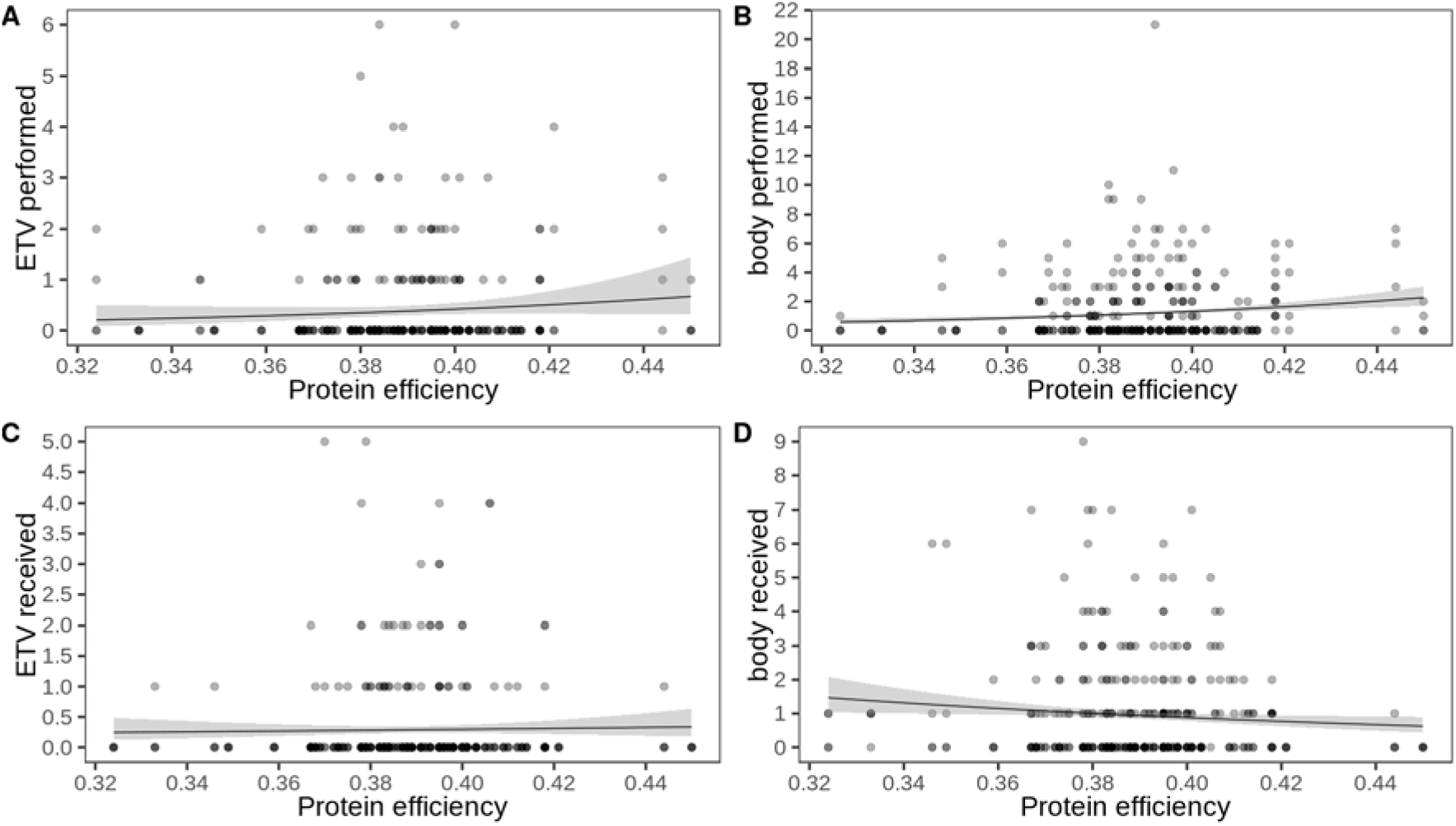
Counts of behaviors directed at or received by pen mates as a function of protein efficiency. **A** Number of problematic behaviors performed, i.e., toward ear, tail, vulva or perineum (ETV), **B** Number of potentially problematic behaviors performed, i.e., toward body or head, **C** Number of problematic behaviors received, i.e., toward ETV, **D** Number of potentially problematic behaviors performed, i.e., toward body or head. Note that the regression line in **A** is quasi-Poisson, for **B** to **D** Poisson (zero-inflation, as well as negative binomial regression lines are not implemented in the function).

### Lesions, tail posture, and cleanliness

Thirteen pigs with a medium-range PE (mean PE = 0.39, min = 0.33, max = 0.42), had minor wounds, and only one pig had a part of the tail missing (PE = 0.41), with a PE slightly above average. The other animals had no wounds or only superficial scratches. Five pigs had a straight tail (mean PE = 0.38, min = 0.33, max = 0.41), and all others had curled tails. We observed one pig having a straight tail over all four observations, another over three, two over two, and one over one observation). No pigs with tucked tails were observed, and all pigs were clean, except one that was slightly soiled (PE = 0.37) during two observations.

### Initiation and outcome of confrontations

The best model for the number of confrontations a pig initiated included PE, but it was not significantly associated, but close to significance (Table 6, Fig. 5). The confidence set contained another model that only included the intercept (Table S3). Only weight was included in the best model for winning a confrontation, but it was not significant (Table 6, Fig. 5). Two of the three other models within a ΔAICc of 2 of the best model indicated that PE and weight are associated with winning. In two of the models, PE was significant, meaning that high PE was significantly correlated with the number of confrontations won. Heavier pigs won confrontations more often. Sex was included in one of the models but was not significantly related to winning (Table S4). The best model for losing a confrontation included only the intercept (Table 6, Fig. 5). The other model in the confidence set included weight, which was not significant (Table S5).

**Table 6:**
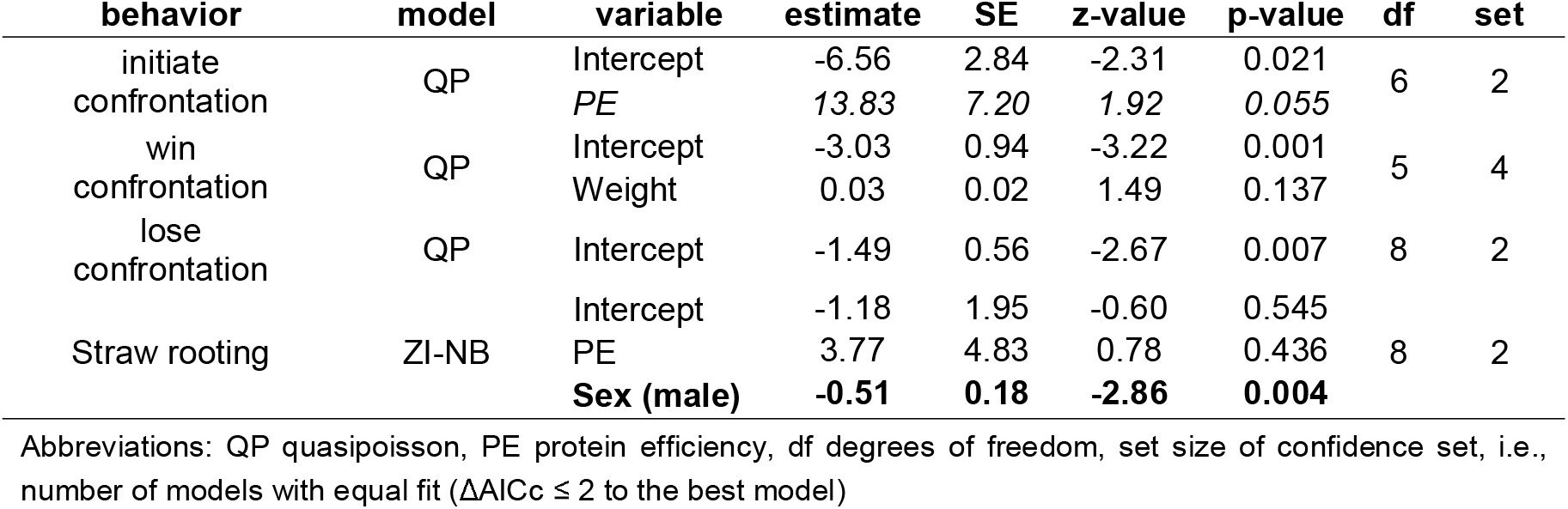
Best models from the model selection for initiation and outcome of confrontations as a function of protein efficiency, sex and body weight and straw rooting as a function of PE and sex. Only the best fitting models in terms of AICc are shown. Significant effects (p < 0.05) are highlighted in bold, and effects with 0.05 ≥ p < 0.10 in italics.

**Fig. 5:**
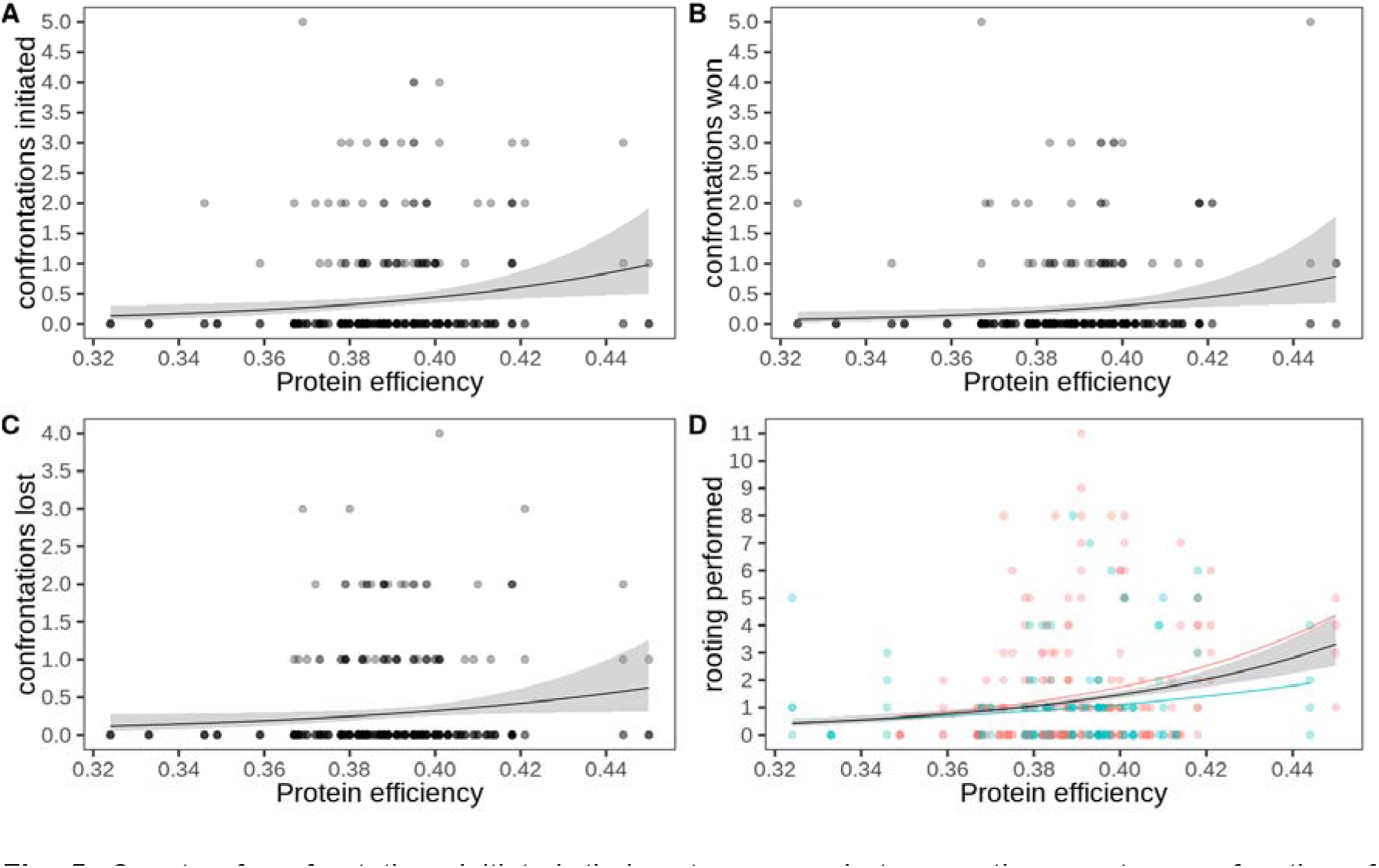
Counts of confrontations initiated, their outcomes and straw rooting counts as a function of protein efficiency. **A** Number of confrontations initiated, **B** number of confrontations won, **C** number of confrontations lost, **D** number of straw rooting bouts performed (females in pink, castrated males in turquoise). Note that the regression line in **A** to **C** is quasi-Poisson, and Poisson in **D** (zero-inflation, as well as negative binomial regression lines are not implemented in the function).

### Straw rooting

The best model for straw rooting included PE, which was not significant, and sex, which was significant (Table 6, Fig. 5). PE was also not significantly associated with straw rooting in the other model with similar fit (ΔACIc ≤ 2 to the best model, Table S6). Sex was significantly associated with the number of straw rooting instances in both models, with females rooting more frequently than castrated males.

## Discussion

The purpose of this study was to investigate the relationship between PE and harmful or potentially problematic behaviors in pigs. The relatively large reduction in dietary crude protein of 20% must be considered when interpreting the results, as this has been associated with an increased prevalence of damaging behaviors. Conversely, the pigs in this study were given fresh, unchopped straw daily, sufficient access to feed, floor space above the legal requirements, and daily pen cleaning, and were closely monitored for signs of damaging behavior. These are all favorable conditions that limit stress and the risk of damaging behavior. Great care was given to formulate a diet that included balanced amounts of EAA.

Deficiencies in EAA, such as methionine, are suspected of fueling tail biting^6, 7^. It should be noted that the experimental diet used in the project represents a relatively strict reduction, which can only be recommended for use in practice if the space available per pig is greater than recommended and the sanitary conditions are optimal, otherwise the risk of tail biting may be increased^6^. Throughout the larger project^19^, from October 2018 to June 2021, when the nearly 700 pigs were slaughtered, piggery staff reported their impression of increased nervousness in the pigs, which they attributed to the protein reduction. One tail biting outbreak was reported in February 2020, before the present behavioral observation study. This outbreak could be contained by providing the pigs with paper bags and hay several times per day. Engaging with these materials, which were novel to them, was sufficient to reduce tail biting. From then on, only 12 instead of 14 pigs were housed per feeding station.

### Damaging or problematic behaviors

In the present study, we did not find evidence that PE is associated with an increased risk of damaging behavior toward pen mates. Only a few pigs showed damaging behavior in terms of biting the ETV regions of pen mates, and these pigs’ PE did not stand out in any direction. Further, there was no evidence that PE made an individual more susceptible to receiving damaging behaviors. However, to assess animal welfare more comprehensively, it is essential to consider other types of behaviors, such as other oral and nasal behaviors, directed at the ETV areas of pen mates (termed “problematic behaviors” in our study), and even behaviors directed at the head and the body (termed “potentially problematic behaviors”), in addition to tail (and ear) biting. Concerning problematic behaviors, we did not find strong evidence of an association with PE, but as PE was present in two confidence set models, we cannot completely rule it out. Furthermore, there was no evidence that PE was linked to potentially problematic behaviors. It is unclear whether the oral and nasal manipulation of conspecifics is harmful, as they represent positive interactions (allogrooming^16, 39^. However, if performed excessively out of boredom or stress, they can affect and stress pen mates^16^ and even lead to injuries.

There was no evidence that sex affected any of the problematic or potentially problematic behaviors, which is consistent with what has been reported previously^8^. However, this might differ when entire males are reared. For instance, Clouard *et al*.^40^ reported that while female and non-castrated male piglets were generally equally active, the latter engaged in more social interactions than females. The increased protein requirements of boars compared to females and castrated animals could also lead to an increased frequency of damaging behavior if the diet does not meet the boars’ needs^12^. It is unsurprising that many injuries occurred outside the 5-min observation periods. Moreover, not all behaviors resulted in visible injuries. Concerning tail posture, which has been suggested as an indicator of tail biting^11^, we could not establish a clear connection to PE, as the five animals in which a straight (hanging) tail posture was observed (all others had raised curled tails) had an approximately average PE. Some pigs with straight tails were observed as being exposed to damaging or potentially problematic behavior, but most straight-tailed pigs were not exposed to an unusual number of behaviors during the observation, therefore it is likely the straight-tail posture indicates interactions that occurred outside the observation intervals. It has to be noted that straight tails are not necessarily linked to tail biting, but also indicate a positive and/or relaxed state in pigs, especially in enriched environments^39^.

The relationship between PE and potentially problematic behaviors presented here is purely phenotypic. However, in the absence of genetic studies, it might be cautiously interpreted as a proxy for genetic correlations^41, 42^. Thus, when engaging in genetic selection for increased PE, a dramatic increase in tail biting due to co-selection is highly unlikely, but it is advisable to monitor pig behavior carefully to detect any deterioration. The total observation time of 20 min per pig in this study is perhaps a rather short period of time in view of the fact that behaviors are labile traits to have any indication of a genetic basis for these traits. The number of behavioral observations available in a study is necessarily limited due to their laborious nature, but several promising computer vision tools are being developed^43, 44^ that may allow automated detection of behaviors that can be used in high-throughput phenotyping for genetic studies.

### Frequency and outcome of confrontations

Using the number of confrontations initiated and their outcomes as an indicator of whether aggression or dominance is associated with efficiency, we found some evidence of the role of PE in how often pigs started a confrontation and how often they won it. While a pig tended to initiate more confrontations when it had a higher PE, which was close to but not significant, there was also evidence that pigs with higher PE won confrontations more often, as PE was included in two of the four models with practically equal fit, and it was significant both times. One might speculate that this could be explained by the greater strength of protein-efficient pigs, since they might have a higher muscle mass. The number of confrontations lost was not linked to PE. If a pen mate occupies a place or resource the focal pig wants to use, there will be a brief confrontation until the matter is settled. Confrontations among familiar pigs in established groups are a normal part of pigs’ behavioral repertoire and reflect the dominance hierarchy in the pen^40^. If the pigs can solve a confrontation rather quickly without causing much injury, these are not a matter of concern.

### Straw rooting

The number of pig-oriented actions was much higher than the number of object-oriented actions (including metal chains, pen barriers, drinking bowls, and straw) in this study, similar to the results of Meer *et al*.^6^, in whose study only a chain with a hard plastic tube was available as enrichment. In our case, straw was provided daily, and the animals engaged with it extensively. However, this might have satisfied only part of their urge to interact with mobile, flexible, and deformable objects^6, 45, 46^. One might speculate that the provision of straw has already prevented some of the abnormal manipulations of pen mates, which would have been more pronounced in the absence of this occupational opportunity^16, 40^. PE did not seem to be linked with the frequency of straw rooting, but we found that females rooted more than castrated males, which has been reported previously^47, 48^.

## Conclusion

The potential consequences of selection for higher PE on pig welfare are thus far unknown, but a trade-off between sustainability and welfare is a concern. In this study, the first to examine the relationship between PE and tail biting or potentially problematic behaviors, we found no evidence of a major risk of an increase in harmful behaviors with higher PE, even in a scenario in which protein in feed is severely restricted. However, the statistical approach used here does not allow us to entirely rule out the possibility that PE may be related to an increased frequency of manipulation of pen mates. While this may not necessarily be problematic, as the behavior was not directed at vulnerable body regions, it could possibly indicate misdirected foraging behavior. Consequently, we believe it is important to provide pigs with an appropriate environment and sufficient space, possibly combined with close monitoring of pens through automatic early warning systems^10^, especially if they are to become more efficient through breeding. Moreover, the effects of breeding for higher PE should be carefully evaluated for possible negative effects on pigs’ physiology and immune systems to avoid these potential causes of damaging behavior. In addition, efforts should also be made to improve the stress susceptibility of pigs through breeding. While this has so far been hampered by low or zero heritability^20^, the development of biomarkers^49,50^and automated phenotyping^43, 44^) could be successful in the future.

## Data availability statement

The data that support the findings of this study are publicly available from Zenodo^51^ (https://zenodo.org/record/5920843), which are described in more detail in Roch *et al*.^25^.

## Acknowledgements

We thank Guy Maïkoff and his team for pig husbandry and Hanno Würbel, Giuseppe Bee, and Catherine Ollagnier for discussions. We are grateful to Madeleine F. Scriba for helpful comments on a version of the manuscript and two anonymous reviewers for their valuable input. This work was financially supported by a grant of the Fondation Sur-la-Croix (https://www.fondation-sur-la-croix.ch/fr) to CK. This study is the result of LR’s master’s thesis, which was awarded the Jean-Pierre Miéville 2021 prize for an outstanding contribution of veterinary medicine to animal welfare.

## Author contributions

Conceptualization: LR and CK. Data curation: LR and CK. Formal analysis: LR, EOE, and CK. Funding acquisition: LR and CK. Investigation: LR and EOE. Methodology: LR and CK. Project administration: LR and CK. Resources: CK. Software: LR, EOE, and CK. Supervision: CK. Validation: CK. Writing – original draft: LR. Writing – review & editing: LR, EOE, and CK.

