## Supplementary information for "Phenotypic link between protein efficiency and pig welfare suggests no apparent trade-offs for mitigating nitrogen pollution"

**Table S1:** Model selection for **A** problematic behaviors (ETV) and **B** potentially problematic behaviors (biting, seizing and manipulation of head and body) performed as a function of protein efficiency and sex. The confidence set, i.e., all models within ΔAICc ≤ 2 from the top model, is separated by a dashed line. Significant effects are highlighted in bold and p-values between 0.05 and 0.10 in italics.

|  | **model** | **variable** | **estimate** | **SE** | **z value** | **p-value** | **df** | **ΔAICc** |
| --- | --- | --- | --- | --- | --- | --- | --- | --- |
| **A** | QP | Intercept | -4.61 | 2.33 | -1.98 | 0.048 | 6 | 0 |
|  |  | PE | 9.31 | 5.95 | 1.57 | 0.117 |  |  |
|  | QP | Intercept | -1.00 | 0.17 | -5.74 | <0.001 | 5 | 0.3 |
|  | NB | Intercept | -4.22 | 2.51 | -1.68 | 0.093 | 6 | 0.8 |
|  |  | PE | 8.36 | 6.43 | 1.30 | 0.194 |  |  |
|  | QP | Intercept | -4.55 | 2.33 | -1.95 | 0.051 | 7 | 1.7 |
|  |  | PE | 9.32 | 5.93 | 1.57 | 0.116 |  |  |
|  |  | Sex (male) | -0.14 | 0.23 | -0.63 | 0.528 |  |  |
|  | ZI-PO | Intercept | 0.12 | 0.19 | 0.66 | 0.512 | 5 | 1.7 |
|  | NB | Intercept | -4.03 | 2.52 | -1.60 | 0.110 | 7 | 2.2 |
|  |  | PE | 8.08 | 6.43 | 1.26 | 0.209 |  |  |
|  |  | Sex (male) | -0.21 | 0.26 | -0.82 | 0.415 |  |  |
|  | ZI-PO | Intercept | -1.60 | 2.12 | -0.76 | 0.449 | 6 | 3.1 |
|  |  | PE | 4.42 | 5.37 | 0.82 | 0.410 |  |  |
|  | ZI-PO | Intercept | -1.38 | 2.15 | -0.64 | 0.522 | 7 | 4.1 |
|  |  | PE | 4.08 | 5.43 | 0.75 | 0.453 |  |  |
|  |  | Sex (male) | -0.25 | 0.24 | -1.05 | 0.294 |  |  |
| **B** | ZI-NB | Intercept | 0.89 | 0.11 | 8.15 | <0.001 | 6 | 0 |
|  | ZI-NB | Intercept | -1.35 | 2.03 | -0.67 | 0.504 | 7 | 0.8 |
|  |  | PE | 5.74 | 5.16 | 1.11 | 0.266 |  |  |
|  | ZI-PO | Intercept | 0.90 | 0.10 | 8.95 | <0.001 | 5 | 2.0 |
|  | ZI-NB | Intercept | -1.35 | 2.03 | -0.66 | 0.507 | 8 | 2.9 |
|  |  | PE | 5.75 | 5.17 | 1.11 | 0.266 |  |  |
|  |  | Sex (male) | -0.03 | 0.20 | -0.16 | 0.874 |  |  |
|  | ZI-PO | Intercept | -1.28 | 2.01 | -0.64 | 0.525 | 6 | 2.9 |
|  |  | PE | 5.57 | 5.11 | 1.09 | 0.276 |  |  |
|  | ZI-PO | Intercept | -1.28 | 2.01 | -0.63 | 0.526 | 7 | 5.0 |
|  |  | PE | 5.58 | 5.12 | 1.09 | 0.276 |  |  |
|  |  | Sex (male) | -0.01 | 0.19 | -0.05 | 0.961 |  |  |
|  | QP | Intercept | 0.00 | 0.15 | -0.03 | 0.974 | 7 | 15.2 |
|  | QP | Intercept | -4.70 | 2.12 | -2.21 | 0.027 | 6 | 16.0 |
|  |  | *PE* | *12.07* | *5.40* | *2.23* | *0.026* |  |  |
|  | QP | Intercept | -4.49 | 2.10 | -2.14 | 0.032 | 5 | 18.8 |
|  |  | *PE* | *11.96* | *5.35* | *2.24* | *0.025* |  |  |
|  |  | Sex (male) | -0.35 | 0.21 | -1.69 | 0.091 |  |  |

Abbreviations: ETV behaviours directed at ears, tails or perineal region of conspecifics, ZI zero-inflated, NB negative binomial, QP quasipoisson, PO poisson, PE protein efficiency, df degrees of freedom, ΔAICc difference in AICc to the best model

**Table S2:** Model selection for **A** problematic behaviors (ETV) and **B** potentially problematic behaviors (biting, seizing and manipulation of head and body) received as a function of protein efficiency and sex. The confidence set, i.e., all models within ΔAICc < 2 from the top model, is separated by a dashed line. Significant effects are highlighted in bold and p-values between 0.05 and 0.10 in italics.

|  | **model** | **variable** | **estimate** | **SE** | **z value** | **p-value** | **df** | **ΔAICc** |
| --- | --- | --- | --- | --- | --- | --- | --- | --- |
| **A** | ZI-PO | Intercept | -0.16 | 0.27 | -0.60 | 0.546 | 5 | 0 |
|  | NB | Intercept | -2.47 | 3.32 | -0.74 | 0.457 | 6 | 3.1 |
|  |  | PE | 3.15 | 8.54 | 0.37 | 0.712 |  |  |
|  | ZI-PO | Intercept | -1.18 | 3.23 | -0.37 | 0.714 | 7 | 3.3 |
|  |  | PE | 2.22 | 8.26 | 0.27 | 0.788 |  |  |
|  |  | Sex (male) | 0.26 | 0.29 | 0.89 | 0.371 |  |  |
| **B** | NB | Intercept | 3.01 | 1.89 | 1.60 | 0.110 | 6 | 0 |
|  |  | PE | -7.90 | 4.85 | -1.63 | 0.103 |  |  |
|  | NB | Intercept | -0.05 | 0.11 | -0.50 | 0.614 | 5 | 0.6 |
|  | NB | Intercept | 2.91 | 1.90 | 1.54 | 0.125 | 7 | 1.7 |
|  |  | PE | -7.77 | 4.87 | -1.60 | 0.111 |  |  |
|  |  | Sex (male) | 0.10 | 0.17 | 0.59 | 0.558 |  |  |
|  | ZI-PO | Intercept | 4.01 | 1.78 | 2.25 | 0.025 | 6 | 25.4 |
|  |  | **PE** | **-9.46** | **4.62** | **-2.05** | **0.041** |  |  |
|  | ZI-QP | Intercept | 4.01 | 1.78 | 2.25 | 0.025 | 6 | 25.4 |
|  |  | **PE** | **-9.46** | **4.62** | **-2.05** | **0.041** |  |  |
|  | ZI-PO | Intercept | 3.89 | 1.80 | 2.16 | 0.030 | 7 | 27.3 |
|  |  | **PE** | **-9.25** | **4.64** | **-1.99** | **0.046** |  |  |
|  |  | Sex (male) | 0.08 | 0.17 | 0.48 | 0.635 |  |  |
|  | ZI-PO | Intercept | 0.35 | 0.12 | 2.77 | 0.006 | 5 | 27.5 |
|  | ZI-QP | Intercept | 0.35 | 0.12 | 2.77 | 0.006 | 5 | 27.5 |

Abbreviations: ETV behaviours directed at ears, tails or perineal region of conspecifics, ZI zero-inflated, NB negative binomial, QP quasipoisson, PO poisson, PE protein efficiency, df degrees of freedom, ΔAICc difference in AICc to the best model

**Table S3:** Model selection for initiation of confrontations as a function of protein efficiency, sex and body weight. The confidence set, i.e., all models within ΔAICc < 2 from the top model, is separated by a dashed line. Significant effects are highlighted in bold and p-values between 0.05 and 0.10 in italics.

| **model** | **variable** | **estimate** | **SE** | **z value** | **p-value** | **df** | **ΔAICc** |
| --- | --- | --- | --- | --- | --- | --- | --- |
| QP | Intercept | -6.56 | 2.84 | -2.31 | 0.021 | 6 | 0 |
|  | *PE* | *13.83* | *7.20* | *1.92* | *0.055* |  |  |
| QP | Intercept | -1.17 | 0.34 | -3.47 | 0.001 | 5 | 1.7 |
| QP | Intercept | -6.74 | 3.15 | -2.14 | 0.033 | 8 | 2.2 |
|  | *PE* | *13.29* | *7.27* | *1.83* | *0.067* |  |  |
|  | Sex (male) | -0.33 | 0.25 | -1.33 | 0.185 |  |  |
|  | Weight | 0.01 | 0.02 | 0.57 | 0.572 |  |  |
| QP | Intercept | -1.33 | 0.93 | -1.43 | 0.153 | 6 | 3.8 |
|  | Weight | 0.00 | 0.02 | 0.19 | 0.852 |  |  |
| NB | Intercept | -6.04 | 3.21 | -1.88 | 0.060 | 8 | 8.6 |
|  | PE | 11.59 | 7.49 | 1.55 | 0.122 |  |  |
|  | Sex (male) | -0.23 | 0.25 | -0.91 | 0.361 |  |  |
|  | Weight | 0.01 | 0.02 | 0.61 | 0.539 |  |  |
| PO | Intercept | -6.13 | 2.99 | -2.05 | 0.040 | 5 | 32.9 |
|  | PE | 12.32 | 7.58 | 1.63 | 0.104 |  |  |
| NB | Intercept | -6.13 | 2.99 | -2.05 | 0.040 | 5 | 32.9 |
|  | PE | 12.32 | 7.58 | 1.63 | 0.104 |  |  |
| PO | Intercept | -1.33 | 0.34 | -3.91 | <0.001 | 4 | 33.6 |
| NB | Intercept | -1.33 | 0.34 | -3.91 | <0.001 | 4 | 33.6 |
| PO | Intercept | -1.21 | 0.87 | -1.39 | 0.164 | 5 | 35.6 |
|  | Weight | 0.00 | 0.02 | -0.16 | 0.876 |  |  |
| NB | Intercept | -1.21 | 0.87 | -1.39 | 0.164 | 5 | 35.6 |
|  | Weight | 0.00 | 0.02 | -0.16 | 0.876 |  |  |
| PO | Intercept | -6.09 | 3.20 | -1.90 | 0.057 | 7 | 36.5 |
|  | PE | 12.12 | 7.60 | 1.60 | 0.111 |  |  |
|  | Sex (male) | -0.19 | 0.26 | -0.72 | 0.470 |  |  |
|  | Weight | 0.00 | 0.02 | 0.14 | 0.886 |  |  |

Abbreviations: QP quasipoisson, PE protein efficiency, df degrees of freedom, ΔAICc difference in AICc to the best model

**Table S4:** Model selection for winning confrontations as a function of protein efficiency, sex, body weight and activity. The confidence set, i.e., all models within ΔAICc < 2 from the top model, is separated by a dashed line. Significant effects are highlighted in bold and p-values between 0.05 and 0.10 in italics.

| **model** | **variable** | **estimate** | **SE** | **z value** | **p-value** | **df** | **ΔAICc** |
| --- | --- | --- | --- | --- | --- | --- | --- |
| QP | Intercept | -3.03 | 0.94 | -3.22 | 0.001 | 5 | 0 |
|  | Weight | 0.03 | 0.02 | 1.49 | 0.137 |  |  |
| QP | Intercept | -1.73 | 0.30 | -5.72 | <0.001 | 4 | 0.1 |
| QP | Intercept | -9.56 | 3.04 | -3.15 | 0.002 | 6 | 1.5 |
|  | **PE** | **20.46** | **7.69** | **2.66** | **0.008** |  |  |
| QP | Intercept | -9.63 | 2.91 | -3.31 | 0.001 | 8 | 1.7 |
|  | **PE** | **16.78** | **7.57** | **2.22** | **0.027** |  |  |
|  | Sex (male) | -0.29 | 0.29 | -0.97 | 0.331 |  |  |
|  | **Weight** | **0.03** | **0.02** | **1.98** | **0.048** |  |  |
| NB | Intercept | -5.97 | 3.14 | -1.90 | 0.057 | 6 | 6.2 |
|  | PE | 11.67 | 8.02 | 1.46 | 0.145 |  |  |
| NB | Intercept | -1.42 | 0.32 | -4.38 | <0.001 | 5 | 6.3 |
| PO | Intercept | -8.86 | 3.28 | -2.70 | 0.007 | 5 | 34.3 |
|  | **PE** | **17.84** | **8.35** | **2.14** | **0.033** |  |  |
| PO | Intercept | -1.89 | 0.31 | -6.10 | <0.001 | 4 | 35.6 |
| PO | Intercept | -9.03 | 3.16 | -2.85 | 0.004 | 7 | 36.2 |
|  | *PE* | *15.49* | *8.13* | *1.91* | *0.057* |  |  |
|  | Sex (male) | -0.27 | 0.33 | -0.81 | 0.419 |  |  |
|  | Weight | 0.03 | 0.02 | 1.45 | 0.146 |  |  |
| PO | Intercept | -3.20 | 0.86 | -3.70 | <0.001 | 5 | 36.6 |
|  | Weight | 0.03 | 0.02 | 1.54 | 0.124 |  |  |

Abbreviations: QP quasipoisson, PE protein efficiency, df degrees of freedom, ΔAICc difference in AICc to the best model

**Table S5:** Model selection for losing confrontations as a function of protein efficiency, sex, body weight and activity. The confidence set, i.e., all models within ΔAICc < 2 from the top model, is separated by a dashed line. Significant effects are highlighted in bold and p-values between 0.05 and 0.10 in italics.

| **model** | **variable** | **estimate** | **SE** | **z value** | **p-value** | **df** | **ΔAICc** |
| --- | --- | --- | --- | --- | --- | --- | --- |
| QP | Intercept | -1.49 | 0.56 | -2.67 | 0.007 | 5 | 0 |
| QP | Intercept | -0.39 | 1.01 | -0.39 | 0.698 | 6 | 0.1 |
|  | Weight | -0.02 | 0.02 | -1.40 | 0.162 |  |  |
| QP | Intercept | -1.57 | 2.61 | -0.60 | 0.547 | 6 | 2.1 |
|  | PE | 0.22 | 6.57 | 0.03 | 0.973 |  |  |
| QP | Intercept | -0.42 | 2.72 | -0.15 | 0.877 | 8 | 3.5 |
|  | PE | -0.11 | 6.53 | -0.02 | 0.987 |  |  |
|  | Sex (male) | -0.20 | 0.24 | -0.86 | 0.392 |  |  |
|  | Weight | -0.02 | 0.02 | -1.16 | 0.245 |  |  |
| NB | Intercept | -1.45 | 0.50 | -2.87 | 0.004 | 5 | 6.7 |
| NB | Intercept | -2.49 | 2.72 | -0.91 | 0.361 | 6 | 8.6 |
|  | PE | 2.68 | 6.88 | 0.39 | 0.697 |  |  |
| PO | Intercept | -1.48 | 0.51 | -2.92 | 0.004 | 4 | 17.1 |
| PO | Intercept | -0.47 | 0.91 | -0.52 | 0.603 | 5 | 17.2 |
|  | Weight | -0.02 | 0.01 | -1.42 | 0.157 |  |  |
| PO | Intercept | -2.21 | 2.42 | -0.92 | 0.360 | 5 | 19.1 |
|  | PE | 1.87 | 6.06 | 0.31 | 0.758 |  |  |
| PO | Intercept | -1.06 | 2.48 | -0.42 | 0.671 | 7 | 21.1 |
|  | PE | 1.41 | 5.89 | 0.24 | 0.811 |  |  |
|  | Sex (male) | -0.09 | 0.21 | -0.45 | 0.656 |  |  |
|  | Weight | -0.02 | 0.02 | -1.28 | 0.202 |  |  |

Abbreviations: QP quasipoisson, PE protein efficiency, df degrees of freedom, ΔAICc difference in AICc to the best model

**Table S6:** Model selection of number of straw rooting bouts as a function of protein efficiency, sex and activity. The confidence set, i.e., all models within ΔAICc < 2 from the top model, is separated by a dashed line. Significant effects are highlighted in bold and p-values between 0.05 and 0.10 in italics.

| **model** | **variable** | **estimate** | **SE** | **z value** | **p-value** | **df** | **ΔAICc** |
| --- | --- | --- | --- | --- | --- | --- | --- |
| ZI-NB | Intercept | -1.18 | 1.95 | -0.60 | 0.545 | 8 | 0 |
|  | PE | 3.77 | 4.83 | 0.78 | 0.436 |  |  |
|  | **Sex (male)** | **-0.51** | **0.18** | **-2.86** | **0.004** |  |  |
| ZI-QP | Intercept | -1.46 | 1.90 | -0.77 | 0.441 | 8 | 1.1 |
|  | PE | 4.59 | 4.72 | 0.97 | 0.331 |  |  |
|  | **Sex (male)** | **-0.51** | **0.17** | **-2.97** | **0.003** |  |  |
| ZI-NB | Intercept | 0.06 | 0.43 | 0.13 | 0.894 | 6 | 4.6 |
| ZI-PO | Intercept | -0.91 | 1.93 | -0.47 | 0.639 | 7 | 5.4 |
|  | PE | 3.25 | 4.82 | 0.68 | 0.499 |  |  |
|  | **Sex (male)** | **-0.51** | **0.18** | **-2.83** | **0.005** |  |  |
| ZI-NB | Intercept | -2.02 | 2.05 | -0.98 | 0.325 | 7 | 5.6 |
|  | PE | 5.32 | 5.14 | 1.03 | 0.301 |  |  |
| ZI-QP | Intercept | 0.09 | 0.40 | 0.23 | 0.821 | 6 | 6.6 |
| ZI-QP | Intercept | -2.23 | 2.00 | -1.12 | 0.263 | 7 | 7.3 |
|  | PE | 5.96 | 5.01 | 1.19 | 0.235 |  |  |
| ZI-PO | Intercept | 0.12 | 0.42 | 0.29 | 0.772 | 5 | 9.6 |
| ZI-PO | Intercept | -1.74 | 2.05 | -0.85 | 0.395 | 6 | 10.8 |
|  | PE | 4.78 | 5.14 | 0.93 | 0.352 |  |  |

Abbreviations: NB negative binomial, QP quasipoisson, PE protein efficiency, df degrees of freedom, ΔAICc difference in AICc to the best model
